# Evolutionary rescue model informs strategies for driving cancer cell populations to extinction

**DOI:** 10.1101/2024.11.26.625315

**Authors:** Amjad Dabi, Joel S. Brown, Robert A. Gatenby, Corbin D. Jones, Daniel R. Schrider

**Affiliations:** Department of Genetics, University of North Carolina, Chapel Hill, North Carolina, USA; Department of Cancer Biology and Evolution, Moffitt Cancer Center, Tampa, FL, USA; Department of Integrated Mathematical Oncology, Moffitt Cancer Center, Tampa, FL, USA; Diagnostic Imaging Department, Moffitt Cancer Center, Tampa, FL, USA; Department of Biology, University of North Carolina, Chapel Hill, North Carolina, USA; Integrative Program for Biological and Genome Sciences, University of North Carolina, Chapel Hill, North Carolina, USA; UNC Lineberger Comprehensive Cancer Center, University of North Carolina, Chapel Hill, North Carolina, USA

## Abstract

Cancers exhibit a remarkable ability to develop resistance to a range of treatments, often resulting in relapse following first-line therapies and significantly worse outcomes for subsequent treatments. While our understanding of the mechanisms and dynamics of the emergence of resistance during cancer therapy continues to advance, questions remain about how to minimize the probability that resistance will evolve, thereby improving long-term patient outcomes. Here, we present an evolutionary simulation model of a clonal population of cells that can acquire resistance mutations to one or more treatments. We leverage this model to examine the efficacy of a two-strike “extinction therapy” protocol, in which two treatments are applied sequentially to first contract the population to a vulnerable state and then push it to extinction, and compare it to a combination therapy protocol. We investigate how factors such as the timing of the switch between the two strikes, the rate of emergence of resistant mutations, the doses of the applied drugs, the presence of cross-resistance, and whether resistance is a binary or a quantitative trait affect the outcome. Our results show that the timing of switching to the second strike has a marked effect on the likelihood of driving the cancer to extinction, and that extinction therapy outperforms combination therapy when cross-resistance is present. We conduct an *in silico* trial that reveals when and why a second strike will succeed or fail. Finally, we demonstrate that our conclusions hold whether we model resistance as a binary trait or as a quantitative, multi-locus trait.

## INTRODUCTION

Cancer is a heterogeneous collection of diseases with a wide variety of outcomes, natural histories and responses to therapy (Fidler 2012; Melo et al. 2013; Chen et al. 2014; Linnekamp et al. 2015; Shoag and Barbieri 2016; Kossaï et al. 2017). Cancerous neoplasms are often comprised of a diverse amalgamation of normal and malignant cells. A defining feature of cancerous cells is that they arise, divide, and proliferate uncontrollably as they adapt to their host environment through natural selection (Brown et al. 2023). Efforts to characterize the clonal evolutionary dynamics of cancer have been ongoing for decades (Farber 1990; Carreira et al. 2014; McFadden et al. 2014; Yu et al. 2014; Bhang et al. 2015; Jamal-Hanjani et al. 2017; Dang et al. 2020; Sandén et al. 2020; Huang et al. 2023). These efforts include experimental investigation of cancer’s cellular kinetics (growth and death rates) and characterization of genetic changes during initial growth and following treatment. Cancer treatments such as immuno– and chemo-therapeutics pose a strong selective pressure on these populations of cells. Ideally, the effects of these drugs on cancer cell populations are strong enough to drive the cancer cells to extinction, but it is common for these populations to evolve resistance to treatment and then resume uncontrolled growth (Wagle et al. 2011; Finn et al. 2012; Awad et al. 2021; Priest et al. 2023). Resistance to therapy represents a major obstacle to long-term progression free survival of patients. A better understanding of the evolutionary processes of cancer cell populations could lead to therapies that better mitigate or even exploit adaptive responses to treatment (Labrie et al. 2022). Modeling studies have proved useful for improving our understanding of the dynamics of cancer initiation, growth, and response to therapy (Cristini et al. 2003; Mallet and De Pillis 2006; Gatenby et al. 2007; Jeon et al. 2010; Sottoriva et al. 2010; Cook et al. 2016; Zangooei and Habibi 2017). Evolutionary models in particular help us understand when and why some treatment strategies will fail, and may inform new strategies (Mumenthaler et al. 2011; Beerenwinkel et al. 2015; Korobeinikov et al. 2017; Zhou et al. 2020; Pressley et al. 2021).

The interactions between a population and selective forces or environmental factors can be investigated using evolutionary and ecological models tailored for cancer, including those that describe the evolution of therapeutic resistance (reviewed in (Merlo et al. 2006) and (Laplane and Maley 2024)). For example, evolutionary rescue models (Gomulkiewicz and Holt 1995; Orr and Unckless 2008; Bell and Gonzalez 2009; Orr and Unckless 2014) define the patterns and dynamics of a population threatened with extinction due to a change in selection forces and the probability that that population will adapt sufficiently to avoid extinction and then subsequently recover. Evolutionary rescue occurs when a population is being driven towards extinction by unfavorable changes in the selective environment, but then a beneficial allele (or combination thereof) that is sufficient to mitigate or even benefit from this change becomes common in the population soon enough that the population can rebound before extinction. This results in a U-shaped trajectory of population size that is the hallmark of evolutionary rescue: initial rapid decline to a nadir, followed by renewed expansion (Orr and Unckless 2014).

Devising cancer treatment strategies with sustained effectiveness is in part tantamount to finding ways to prevent evolutionary rescue. In cancer, the population consists of the tumor cells, the selective force driving the population towards extinction is the therapy, and the beneficial allele is the mutation that confers drug resistance. Whether a tumor cell population in decline due to treatment will experience rescue depends on the chance that a resistant clonal subpopulation is established soon enough to prevent extinction. This in turn depends on the size of the cancer cell population at the onset of treatment, the rate at which the treatment kills cancer cells (which together determine how long it will take the cancer population to go extinct unless rescue occurs), and the mutation rate for resistance alleles. Higher mutation rates increase the probability that rescue will occur at any point in time. The presence of preexisting resistance mutations also increases the probability of evolutionary rescue (Orr and Unckless 2008). Thus, preventing evolutionary rescue demands therapeutic regimes that maximize the selection against non-resistant tumor cells, while minimizing or delaying mutational events that give rise to resistance. For example, evolutionary rescue theory implies that combining multiple therapeutic drugs with independent modes of action can greatly improve patient outcomes because the rate of appearance of cells resistant to each drug in combination would typically be far lower than the mutation rate for resistance to a single drug. This notion is supported by the success of combination therapy in comparison to single therapy protocols (Labrie et al. 1993; Li et al. 2014; Mokhtari et al. 2017; Palmer and Sorger 2017; Fisusi and Akala 2019; Plana et al. 2022).

An alternative approach is “extinction therapy” (Gatenby et al. 2019; Gatenby et al. 2020; Gatenby and Brown 2020; Felder et al. 2021). Extinction therapy proposes that after a radical change in the environment resulting in a population crash, the population could be driven to extinction by a comparatively minor environmental perturbation that otherwise would not be a major threat. At the onset, extinction therapy resembles standard-of-care treatment: the most effective available therapy is initially used, continuing this treatment as long as it remains effective. Once the cancer cell population size has been dramatically reduced by the first-line treatment, a “second strike” is introduced with a different drug that may be less effective but is sufficient to push the now-threatened cancer cell population to extinction. Evolutionary modeling has shown that this “two-strike” extinction therapy can be effective (Gatenby et al. 2020; Patil et al. 2023), presuming the second strike is not too early (before the population has shrunk enough to be threatened by a less-effective treatment) or too late (once the population has been rescued by resistance to the first treatment and recovered to a large enough size to survive the second treatment).

Evolutionary modeling can answer outstanding fundamental questions about the conditions under which extinction therapy would be a superior approach, and how it should be carried out. Here we ask: how do factors like the mutation rate and the rate at which a therapy kills cancer cells influence the probability that a two-strike treatment will achieve extinction? What is the optimal timing of the second strike? Perhaps most importantly, when will simultaneous combination therapy be more effective than a sequential two-strike therapy, and vice versa?

Using population genetic simulations, we investigate the optimal timing of the second strike in an extinction therapy treatment protocol and compare the probability of extinction achieved by this sequential two-strike therapy with that of combination therapy. Our simulation approach reveals: 1) interactions between the optimal timing of two-strike sequential therapy, the mutation rate, and the rate at which the drug kills cancer cells; 2) the scenarios in which combination therapy outperforms optimally timed sequential extinction therapy and vice-versa, and 3) the implications of the genetic architecture of resistance (i.e., a binary trait caused by a single mutation vs. a quantitative trait resulting from the cumulative effect of many mutations) for the optimal timing and effectiveness of a second strike.

## METHODS

Several approaches have been used to model the growth, response to therapy, and clonal evolution of cancer. These include mathematical models (Dingli et al. 2006; Benzekry et al. 2014; Sun and Hu 2018; Zhang et al. 2022), spatial models (Thalhauser et al. 2010; Komarova 2013; Waclaw et al. 2015), and agent-based models (Wang et al. 2007; Zhang et al. 2009; Mustapha et al. 2016; Gong et al. 2017).

We used the software `SLiM 4.0` (Haller and Messer 2023) to simulate a diploid clonal population of cells undergoing expansion, treatment with two drugs, and mutations that confer resistance to these drugs. This software allows for agent-based evolutionary simulations which are widely used in the field of population genetics and provide a great deal of flexibility in modeling the evolution of resistance with minimal mathematical assumptions. Cells in these simulations have a specified genome size in base pairs and the simulation has a set mutation rate per base pair per cell cycle. To model clonal evolutionary dynamics, we set the recombination rate to 0. Events in the simulation occur over discrete generations (called cycles in SLiM), with the following events occurring sequentially during each generation (Figure S1):

1. Cell replication: Each cell initially undergoes mitotic replication, producing two daughter cells with the exact same genome (i.e. containing the same set of mutations) as the parent cell. Thus, the cancer cell population size doubles during this step.
2. Mutation: After cell division, each daughter cell may acquire new mutations that affect resistance to treatment (see step 3). Mutations can randomly occur at sites along the genome according to the mutation rate per base pair per cell cycle. If a mutation occurs, its type (e.g. conferring resistance to drug 1 vs. conferring resistance to drug 2) is determined according to a discrete probability distribution of mutation types, referred to as the draw rates. For instance, if the draw rates of two mutation types are (0.5, 0.5), then each mutation type is equally likely to be chosen.
3. Calculation of fitness costs: This step involves tallying the number of mutations in a cell and then calculating its fitness depending on the presence or absence of treatment during the current generation. Treatment reduces fitness, but resistance mutations can mitigate or eliminate this fitness cost. Cells may also die in the absence of treatment (“turnover”).
4. Death and survival: Next, the fitness costs incurred by each cell are used to calculate the probability of survival. At this step, each cell has a probability of survival, *P_surv_*, while the remaining cells die and thus removed from the simulation. Thus, the population contracts during this step, and cells that survive return to step one (replication) in the next cell cycle.

### First model: two-strike extinction therapy in an evolutionary rescue model

A simulation begins with the expansion phase: a population starting with a single cell grows exponentially as described above until reaching a size of ∼2^%&^. We note that this population size of about one million cells is smaller than the number of cells found in a clinically relevant tumor, but simulating smaller populations is a useful way of modeling the dynamics of larger populations while maintaining computational efficiency (but see (Dabi and Schrider 2024). Next comes the treatment phase, which consists of the application of the first drug for a specified number of time steps, followed by immediately switching to the second drug. The number of cell cycles during which the first drug is applied is referred to as the “second-strike lag time”. Throughout the simulation, two types of mutations can occur: m1 and m2, which confer resistance to the first and second drugs, respectively. However, these mutations confer no fitness advantage or cost during the expansion and thus evolve under purely random drift throughout this phase. Note that we did not allow back-mutations: once a cell has acquired a given resistance mutation, all its descendants will harbor that same resistance allele.

In this model, resistance is a binary trait, and resistant cells do not experience any reduction in survival due to treatment nor due to any fitness tradeoff associated with resistance. The magnitude of the effect of each drug on the survival of non-resistant cells (“dose”) during its treatment application is a tunable parameter. The equation governing survival probability during step 4 of each generation is as follows:

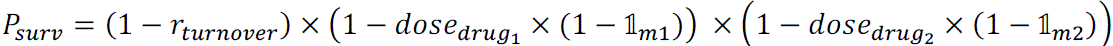

where:

*r_turnover_* is the turnover rate of cells. We set this to 0.20 for all simulations (i.e. 80% of cells survive in the absence of treatment)

*dose_drug_*_1_ is the dose of the first drug,

*dose_drug_*_2_ is the dose of the second drug,

𝟙*_m_*_1_ is the indicator function that the cell has a mutation resistant to drug 1, and 𝟙*_m_*_2_ is the indicator function that the cell has a mutation resistant to drug 2.

During the expansion phase, both drug doses are zero, meaning fitness is only governed by the turnover rate. Application of either drug entails setting its dose to a specified value > 0 while the other drug’s dose remains at zero. The effect of each drug’s dose is to reduce the probability of survival, and the indicator resistance functions govern whether a cell will experience this reduction in survival probability. Note that we use the shorthand “drug” to refer to each treatment, but each of these treatments could themselves be thought of as a combination of multiple drugs that together kills cancer cells according to the specified dose parameter and to which resistance evolves at the specified mutation rate.

We note that this simulation model differs from that of (Gatenby et al. 2020) extinction therapy simulations in several key respects: 1) We model the two strikes in the same manner in that they are continually applied drugs that each reduce the fitness of cancer cells, while Gatenby et al. modeled the second strike as an instantaneous 40% reduction in the cancer cell population size rather than a continually applied treatment; 2) We end treatment with the first drug at the moment the second strike begins, whereas Gatenby et al.’s model continues therapy 1 after the second strike 3) Gatenby et al. modeled resistance as a quantitative rather than binary trait (a change we experiment with below); and 4) Gatenby et al.’s model includes an Allee effect, while ours does not.

For our first set of simulations, we varied the mutation rate and second-strike lag time to test the effect these parameters on treatment success. We tested per-base pair mutation rates between 1.0 × 10^-6^ and 3.1 × 10^-5^ with increments of 2 × 10^-6^ and second-strike lags between 1 and 20 cell cycles with increments of 1 cell cycle. Our drug dose effects in these simulations were fixed at 0.75 for both drugs (i.e. 75% of sensitive cells are killed by the drug each generation, on average). We ran 100 replicates of each parameter combination and measured the fraction of replicates in which the population went extinct, which we refer to as the “fraction of extinct replicates.” We also tested the effect of using a lower dose of either drug across the various mutation rates. Furthermore, we fixed the mutation rate and examined the effect of varying both drug doses simultaneously.

### Fixed trajectories simulations

In addition to examining sets of independent simulation replicates, we also conducted an *in silico* trial examining how outcomes of two-strike extinction therapy differed among a set of 55 *in silico* patients, where each patient is defined by a combination of mutation rate and drug dose. Specifically, under a variant of our first model, we examined the dynamics leading to extinction or evolutionary rescue for each patient by fixing the trajectory of the population up to the time of applying the second treatment—this was achieved by using a fixed initial random seed that was unique to that patient. Therefore, all replicates of patient share not only the drug dose and mutation rate, but the entire population trajectory up until the second treatment, including all birth and death events and mutations. When the second strike happens, a new replicate-specific random seed is chosen, producing stochastic trajectories after the second strike and allowing us to examine the distribution of treatment outcomes for each lag time for each synthetic patient.

### Second model: the effect of combination therapy

We compared a combination therapy consisting of the simultaneous application of both drugs to sequential two-strike extinction therapy, again considering a range of second-strike lag times for the latter. In these simulations, combination therapy is simulated as both drugs having constant doses higher than zero throughout the treatment phase. The dose effects of both drugs in these simulations are always equal, and the lowest dose used for each drug in the combination was 0.4. We note that when both drug doses are 0.4, the survival probability of cells is still below 0.5 (see survival probability formula in Methods), meaning that a net decrease in population size is expected in the absence of resistance. For the two-strike simulations included in this comparison, we fix the dose effect of both drugs to 0.9, while varying this dose effect during combination therapy. We ran 100 replicates of each parameter combination.

### Third model: cross-resistance

We modified the second model to include a third rare mutation type which confers resistance to both drugs. The draw rates for m1, m2, and m3 mutations under this third model are 45%, 45%, and 10% respectively. The survival probability under this model is:

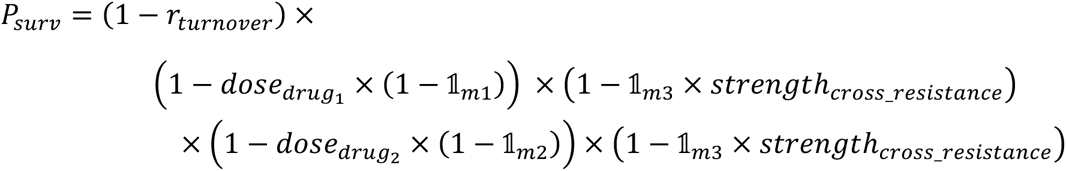

This equation shows that the presence of cross-resistance mutations (i.e. 𝟙*_m_*_2_ = 1) dampens the effect of both drugs on fitness in proportion to the strength of the cross-resistance. Here, a cell that lacks a mutation of type m1 or m2 can still be resistant to both drugs if it has a mutation of type m3, but rather than full resistance, the degree of resistance imparted by this mutation is determined by *strength_cross_resistance_*; if *strength_cross_resistance_* = 0 then m3 provides no resistance, but if *strength_cross_resistance_* = 1 then m3 confers full resistance to both drugs. We ran these simulations across a range of values of *strength_cross_resistance_*, comparing the outcomes of combination therapy to sequential therapy with various lag times.

### Fourth model: varying drug doses and the mutation rate ratio in combination therapy

To investigate whether the optimal combination of doses during combination therapy depends on the relative rates of mutation to resistance alleles for the two drugs, we carried out simulations with varying ratios of the rates of m1 and m2 mutations (achieved by varying the rate of m1 mutations while fixing the rate of m2 mutations). Specifically, we let the mutation rate for m1 mutations rage from 1 × 10^-5^ to 1.01 × *e*^-3^ while the rate for m2 mutations was fixed to 1.5 × 10^-5^. We also varied the ratio of the doses of the two drugs, under the constraint that (1 – *dose_drug_*_1_) × (1 – *dose_drug_*_2_) is fixed at 0.25. This was done by setting *dose_drug_*_1_ to a value between 0.30 and 0.70 and then setting *dose_drug_*_2_ as follows:

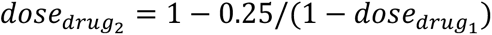

This ensures that the probability of survival for non-resistant cells is the same across all parameter combinations for this model. Any differences in extinction probability across parameterizations are therefore due to the interplay between drug mutation rates and drug doses.

### Simulations modeling resistance as a quantitative trait

We also experimented with a model where drug resistance is a quantitative trait (Gomulkiewicz and Holt 1995). Cells can accumulate mutations conferring resistance to either drug. Each mutation contributes additively to resistance against the corresponding drug and its contribution is drawn from a lognormal distribution. The parameters of this distribution were chosen so that the average resistance effect of a new mutation was 0.1, meaning that, on average, 10 accumulated mutations of a specific type should confer full resistance to the corresponding drug. This yielded a mean of –2.42759 and a standard deviation of 0.5. The survival equation then becomes:

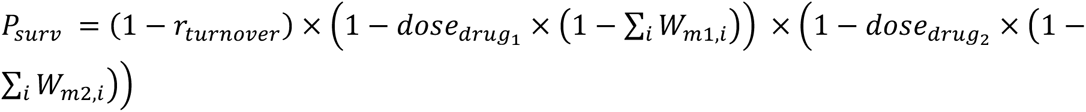

where:

*r_turnover_*, *dose_drug_*_1_, and *dose_drug_*_2_ are as defined above, *W_m_*_1,*i*_ is the fitness effect of the m1 mutation at *i* (zero if no such mutation present), and *W_m_*_2,*i*_ is the fitness effect of the m1 mutation at *i*, if present. Thus, ∑*_i_ W_m_*_1,*i*_ and ∑*_i_ W_m_*_2,*i*_ represent the cell’s total amount of resistance to drugs 1 and 2, respectively

Note that under this model mutations can only increase resistance, and the combined effect of all resistance mutations in a genome was capped at 1.

## RESULTS

### Progression after treatment is a case of evolutionary rescue

Typical cancer treatment involves applying the maximum tolerated dose (MTD) of a first-line therapy until either the cancer is eradicated or until progression is observed (Kareva et al. 2015). Progression often occurs because of the emergence of treatment-resistant cells within the cancer cell population, which quickly propagate until the remaining cells in the population are resistant to the treatment rendering it ineffective. This is an example of evolutionary rescue, wherein a population is rapidly crashing because of a change in the selective environment (i.e. treatment), but one or more adaptive (e.g. resistance-conferring) mutations become common enough in the population prior to extinction, thereby allowing the population to recover and eventually progress beyond the size observed before treatment. A typical example of the trajectory of a population undergoing evolutionary rescue is shown in Figure 1. In this example, the population size is initially growing exponentially, but then rapidly declines following the application of treatment. Eventually, the population size decreases to such a low level that it may be undetectable. However, when the population is approaching its nadir, a subpopulation of resistant cells emerges and begins to become more prevalent, ensuring that the treatment will not succeed in eradicating the cancer.

**Figure 1.**
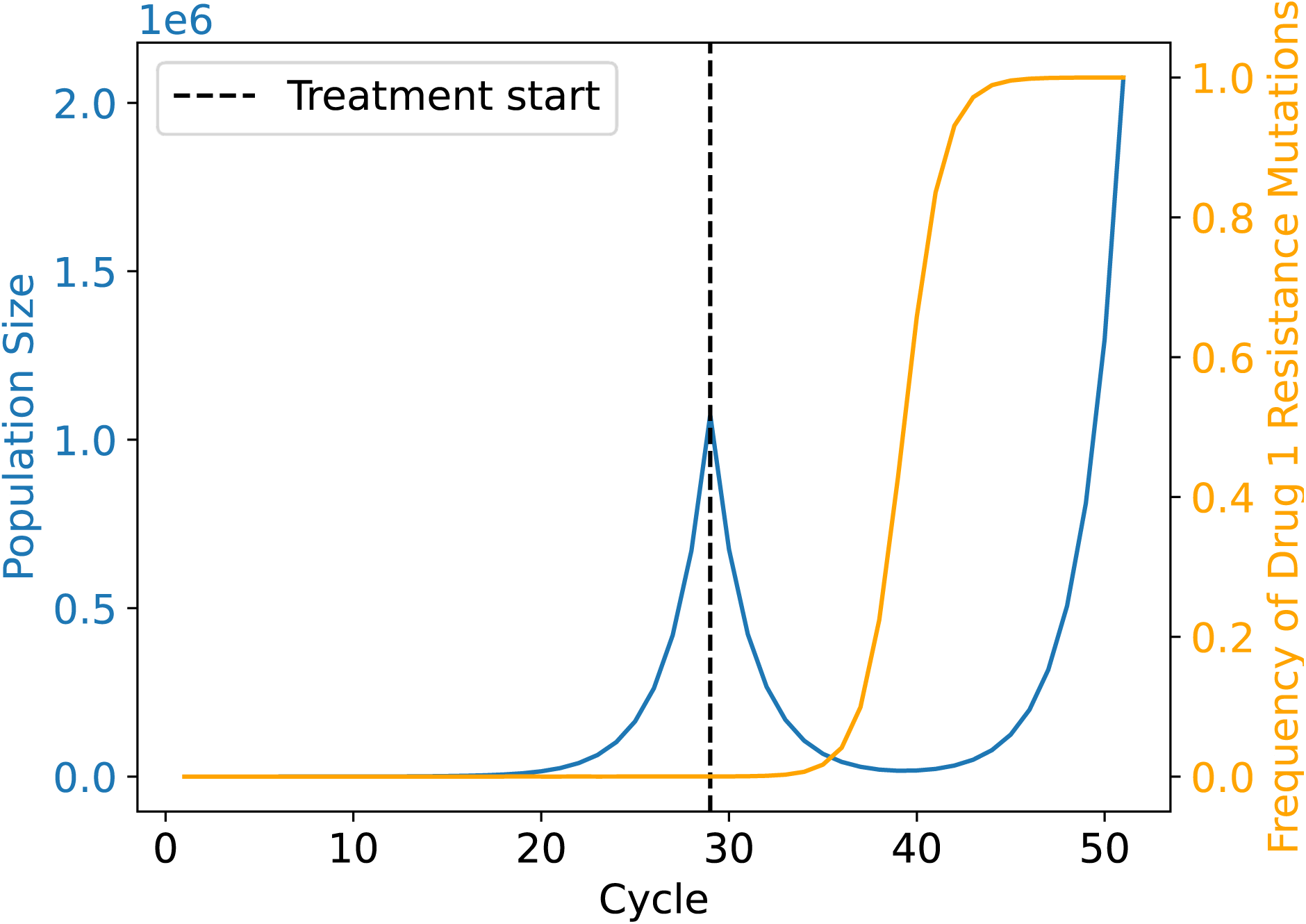
Trajectory of a cancer cell population during evolutionary rescue following treatment. The population expands exponentially (blue line) until the first strike therapy is applied (dashed black line), after which the population declines rapidly. However, as the population is declining, a resistance mutation has emerged and begun to increase in frequency in the population. As the treatment continues, the frequency of the resistance mutation continues to increase (orange line), the population eventually becomes fully resistant, resumes its exponential expansion, and ultimately recovers and progresses beyond its initial size pre-treatment. This is the classic U-shaped curve of evolutionary rescue. This example trajectory is taken from a single simulation replicate of our first model (Methods), but without the application of a second strike. For this replicate the mutation rate was set to 1.5 × 10^-5^ and the drug dose to ∼0.61 (indicating that the fitness of treatment-sensitive cells is multiplied by ∼0.39 during treatment; see Methods).

Differences in treatment regime can alter the trajectory of evolutionary rescue. Below we assess the efficacy of alternative treatment strategies that are designed to reduce the probability of evolutionary rescue of the cancer cell population.

### The optimal second-strike lag is independent of the mutation rate

In a two-strike cancer treatment scenario the timing of the change from the first-line to the second-line therapy may profoundly effect treatment success (Gatenby et al. 2020). We used simulations to estimate the probability of extinction across different timings of the second strike (the “second-strike lag time”) and different rates of mutation to treatment-resistance alleles to understand the relationship between second-strike lag time and the probability that the cancer cell population will be driven to extinction. Our results (Figure 2A) show that there exists an optimal lag time between the onset of the first and second strikes for maximizing the extinction probability of the cell population. In general, the probability of extinction is low when the second drug is applied immediately following the application of the first (a lag time of only 1 cell cycle), then increases with greater lag times until an optimum is reached, and then decreases with increasing lag time. While there is no clear relationship between the mutation rate and the optimal lag time (Figure 2B), which remains generally between 7 and 9 cell cycles in our model, the mutation rate does exert an effect on the dynamics of population extinction probabilities across lag times. For instance, at lower mutation rates the extinction probability exhibits a broad plateau around the optimal lag time, implying that missing the optimal lag time is somewhat forgiving when resistance alleles do not frequently arise. As the mutation rate increases, however, the difference in extinction probability between the optimal lag time and suboptimal becomes starker, with a sharp peak around the optimum, and the extinction probability is also lower in general for all lag times.

**Figure 2.**
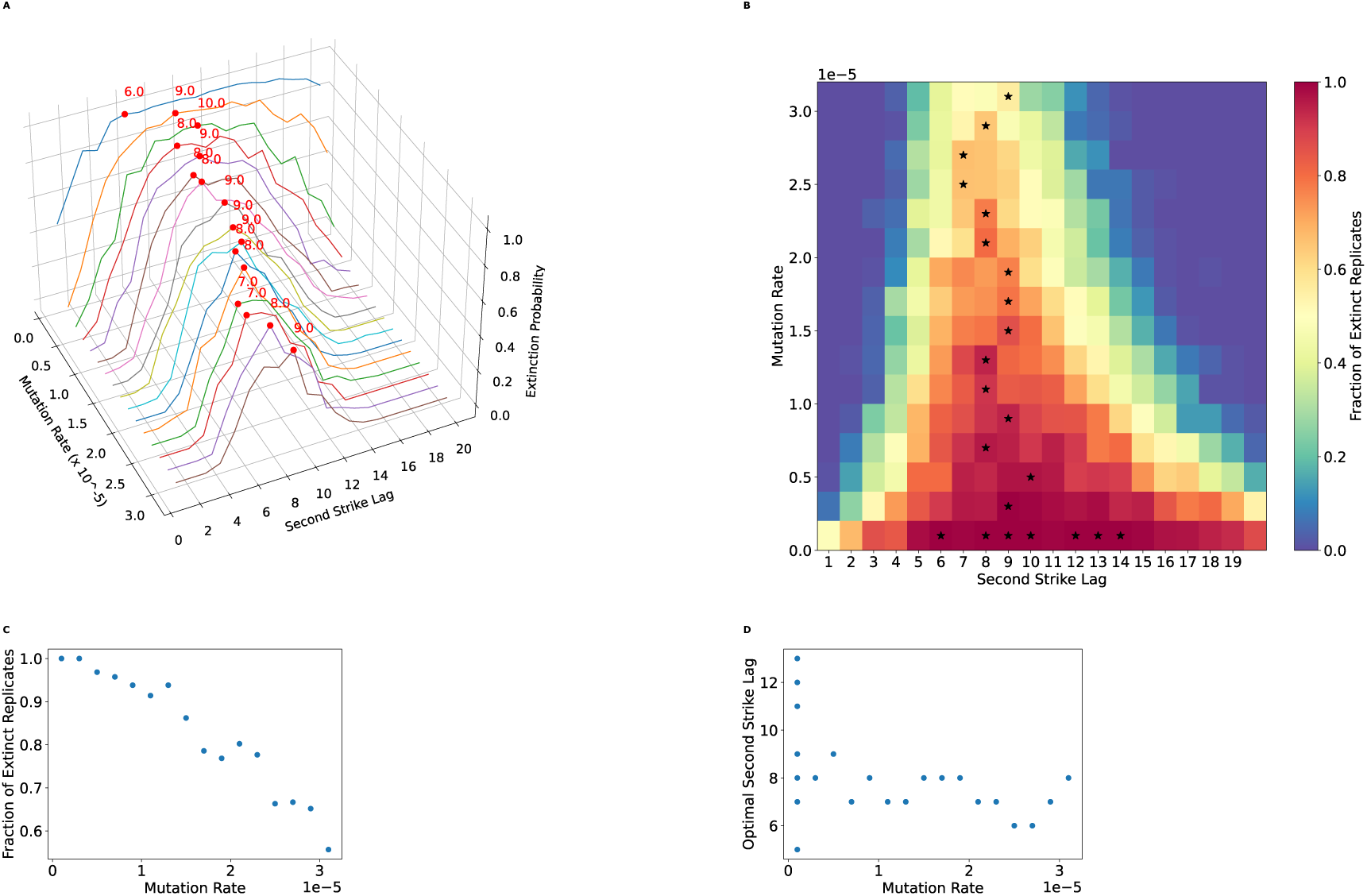
The optimal lag time is unaffected by resistance mutation rate, but the second strike is effective across a wider window of lag times when the mutation rate is lower. A) Extinction rate vs mutation rate and lag time. The *x*-axis represents the per base pair mutation rate of resistant mutations. The *y*-axis (Second Strike Lag) represents the time spent under the first treatment before switching to the second treatment. The *z*-axis (Extinction Probability) represents the fraction of replicates where the population went extinct. Red dots indicate the optimal lag time for each mutation rate (i.e. the lag time with the highest fraction of extinct replicates). The optimal lag time is relatively invariant across mutation rates. However, as the mutation rate increases, the difference in the extinction probability between the optimal lag time and other lag times becomes starker, and the extinction probability decreases for all lag times. B) The window for optimal strike is wider at lower mutations rates. Color indicates the extinction probability estimate and black stars at the optimal lag time(s) for each mutation rate—multiple stars are drawn in the case of a tie. C) The per-base pair mutation rate vs the extinction probability at the optimal lag time for that mutation rate. As the resistance mutation rate increases, the extinction probability decreases. D) Optimal second-strike lag vs mutation rate. This represents a 2D projection of the points indicated by the red dots in A. Despite some stochasticity present, the mutation rate is uncoupled from the optimal second-strike lag time. However, the window for the optimal time is wider for lower mutation rates (panel B), as multiple second-strike lag times produce a high extinction rate for the lower mutation rate.

### More effective therapeutics should be used first during extinction therapy

The rate at which a population is being driven to extinction is a key determinant of the probability and timing of evolutionary rescue (Orr and Unckless 2008, 2014). We examined the effect of our drug dose parameters on extinction probabilities and optimal lag times by testing a scenario with a lower dose of drug 1 than drug 2, and one with a lower dose of drug 2 than drug 1. In this context, drug doses control the reduction in fitness for non-resistant cells during the application of the drug (see Methods for fitness equation). When the first drug has a dose of 0.65 and the second a dose of 0.75, the optimal lag time shifts to around 10 generations (Figure 3A), versus 7–9 generations in the equal-dose effect scenario shown in Figure 2. This increase is likely due to the increased time required to shrink the population of cells enough for the second strike to result in extinction. Again, the mutation rate appears to have little effect on this optimal lag time, and extinction probabilities are lower for all mutation rates compared to the case where both drugs have a dose of 0.75.

**Figure 3.**
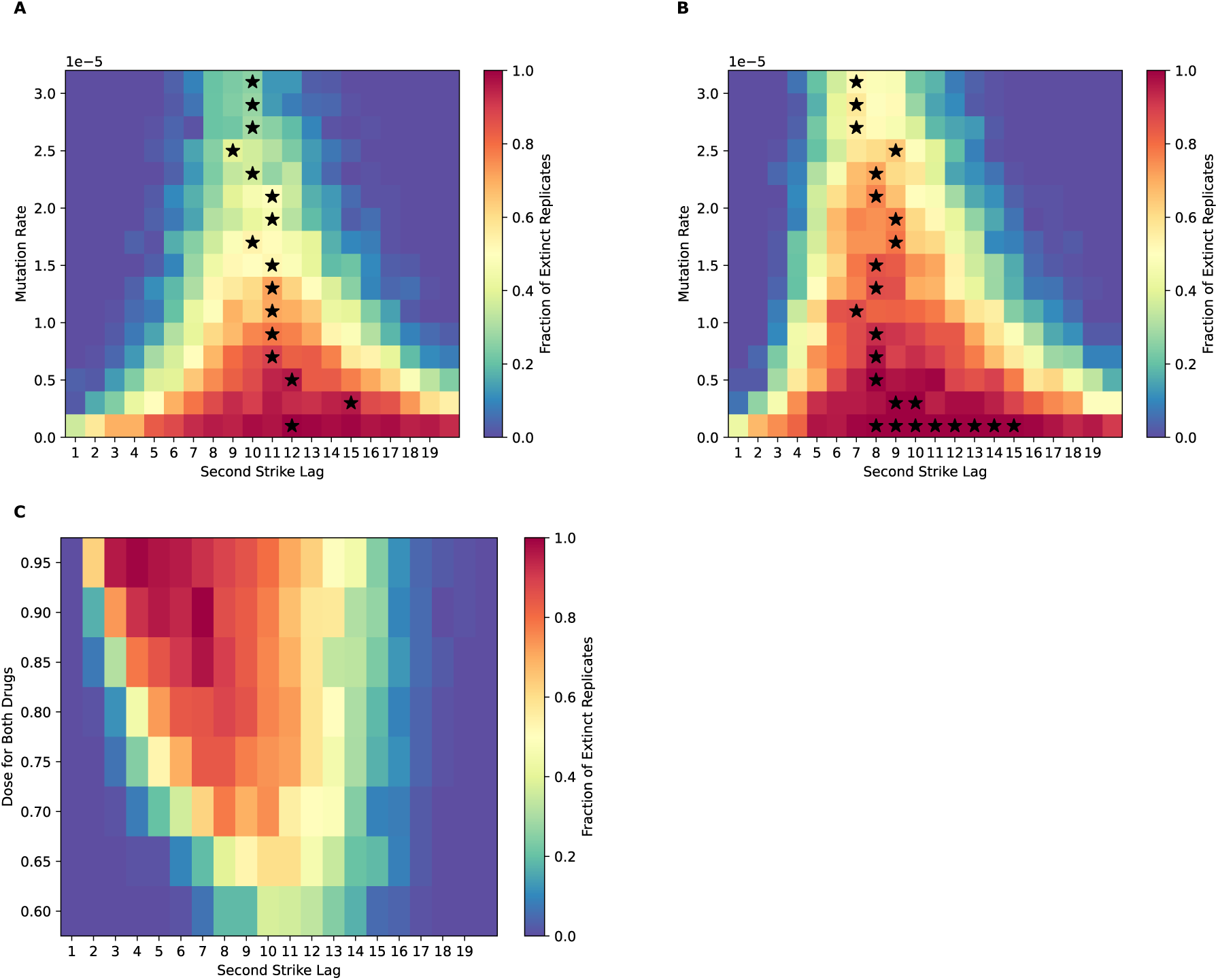
Using a lower-dose drug first reduces overall treatment success and shifts the optimal lag time higher. A) Extinction probability vs mutation rate and lag time when the first drug has a lower dose (0.65) before switching to a second drug with a higher dose (0.75). B) Extinction probability vs mutation rate and lag time when the first drug has a higher dose (0.75) followed by treatment with a second drug with a lower dose (0.65). D) Extinction probability vs lag time and the dose of both drugs when the mutation rate is fixed at 1.5 × 10^-5^, with the dose always being equal for both drugs. Black stars represent the optimal lag time(s) at each mutation rate.

Next, we examined the case where the first drug has a higher dose (0.75) than the second (0.65)—a scenario is more relevant to standard-of-care and to current thinking about extinction therapy (Gatenby et al. 2019) that advocates for using the drug with the highest kill rate as the first-line treatment. We found that optimal lag times in this scenario resemble those for the case in which both drugs had an equal dose of 0.75 and are also not dependent on mutation rate. However, the extinction probabilities across all mutation rates are higher compared to the case of the first drug having the lower dose (Figure 3B vs Figure 3A). For instance, at a mutation rate of 1.5 × 10^-5^, simulations where both drugs have a dose of 0.75 result in an extinction probability of ∼86% at an optimal lag time of 8, but this decreases to 0.53% when the first drug has a lower dose effect (in this case, at an optimal lag time of 11). On the other hand, reducing the dose effect of the second drug but not the first has hardly any effect on the outcome (extinction probability of ∼85%, again with an optimal lag of 8). It is likely that the higher optimal lag time necessary to sufficiently reduce the population size when the first drug has a weaker effect also results in a higher likelihood of resistant mutants appearing. Thus, our results suggest using the stronger drug as the first strike during extinction therapy provides the best chance of eradicating the cancer cell population.

Finally, we set both drugs to an equal dose and examined extinction dynamics across various dose values (with the mutation rate set to 1.5 × 10^-5^ in each instance). When the dose of both drugs is equal, the optimal time increases as the drug dose decreases, and the extinction probability decreases from ∼100% at a drug dose of 0.95 to ∼20% at a drug dose of 0.60 (Figure 3C).

### Investigating the dynamics of evolutionary rescue in extinction therapy via *in silico* trials

#### Extinction therapy succeeds when the first strike removes variants that confer resistance to the second strike

Our simulation approach allows us to go a step further and ask, for a given patient’s disease course, what would have been the optimal treatment strategy? Such information would not only improve our understanding of the dynamics impacting the probability of extinction, but also help us assess the potential of treatment strategies tailored to individual patients. To this end, we conducted an *in silico* trail with a set of patients receiving two-strike extinction therapy, and determined, for each specific patient, the optimal lag time. This may differ from patient to patient both because of different parameter values (e.g. the mutation rate and drug dose parameters differ across patients) and because of stochasticity in the timing of the emergence of resistance and other random events occurring both before and during the treatment course.

Our trial consisted of a set of 50 *in silico* patients, each of whom experienced a deterministic disease trajectory up until the start of the second strike, at which point we performed a set of 100 stochastic simulations to observe that patient’s distribution of outcomes for a given lag time (Methods). Illustrative example patients are shown in Figure 4; the remainder are available on our project’s GitHub repository. An examination of these results reveals that two different, but related factors govern the probability of treatment success (i.e. the fraction of extinct replicates) for a given lag time. The first is the size of the cancer cell population at the time of the second strike, and second is the frequency of the mutations at that time—results that are consistent with analytical results (Orr and Unckless 2008). We therefore might expect the optimal lag time to be the moment the population size is at its minimum (*t_min_*), and indeed in many cases we see that the extinction probability peaks when the second strike begins at *t_min_*. Two such examples are shown in Figure 4C and 4D; for these two synthetic patients a plateau of extinction probability begins when the second strike starts at *t_min_*.

**Figure 4.**
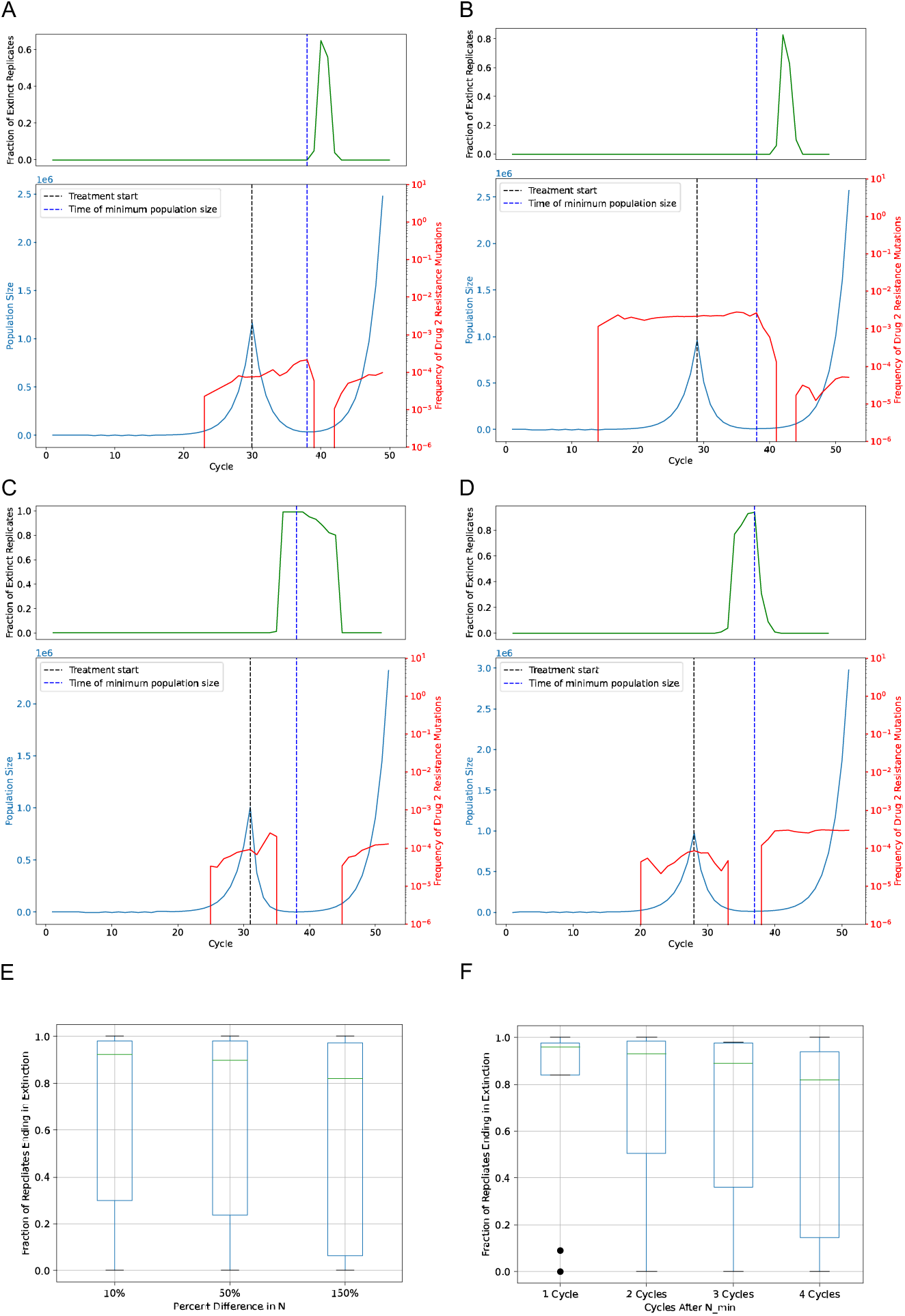
Synthetic patient trajectories show the success of strategies that involve switching treatments near the population nadir and the importance of eliminating standing variation. A-B) Trajectories of synthetic patients where the optimal lag time occurs after the time of minimum population size (dashed blue vertical line) due to remaining standing variation that confers resistance to the second treatment. C-D) Trajectories of synthetic patients where the optimal lag time is at the population minimum because standing variation that confers resistance to the second treatment has been eliminated prior to the population reaching its minimum. Blue lines represent the population size, red line represent the frequency of the mutation conferring resistance to the second strike, and green lines in the upper sub-panels represent the fraction of replicates that have undergone extinction when switching to the second strike at that time. Results for the remaining 46 in silico patients are available on our GitHub repository. E) The distribution of extinction probabilities for a separate set of validation patients across three different timing strategies for the second strike. Each value on the *x*-axis shows the percent growth in population size over the size of the minimum population at the time of switching from the first strike to the second strike. F) The distribution of extinction probabilities for the set of validation patients across three more timing strategies for the second strike; each value on the *x*-axis shows the number of cycles allowed to occur after reaching the minimum population during the first strike before applying the second strike.

However, beginning the second strike at *t_min_* is not the optimal strategy for all synthetic patients. This is because in some cases there is standing variation for resistance to the second treatment present at *t_min_*. Although they confer no fitness benefit prior to the second strike, resistance mutations for the second drug typically emerge during the expansion phase and persist at low frequency, and sometimes remain present even as the population declines to its minimum size (see Figures 4A and 4B for two such examples). In such cases, beginning the second strike at *t_min_* will almost always fail to achieve extinction. Instead, continuing the first treatment for one or more cycles beyond *t_min_* may reduce the frequency of these resistance mutations or even remove them from the population, resulting in a much higher extinction probability. This makes sense because, during the first strike, cells that are resistant to the first treatment are selected for at the expense of others, including those that have preexisting resistance to the second treatment, thereby reducing the chance of evolutionary rescue following the second strike (e.g. Figures 4A, B).

#### In silico *trials inform strategies for timing the second strike*

Our synthetic trials also identify two-strike treatment strategies that are more likely to prevent evolutionary rescue of the cancer cell population. For example, when examining the results for all 50 synthetic patients, we find that switching to the second treatment at the time where the population is at or near its minimum after the first strike is generally a successful strategy (e.g. when the cancer cells’ mutation rate is set to 6 × 10^/0^, the strategy of beginning the second strike at *t_min_* results in an extinction probability that is within 3% of its maximum for ∼70% of synthetic patients).

In practice, however, it may not be possible to known *t_min_*, until after the population has begun to rebound. We therefore assessed the effectiveness of beginning the second strike after some amount of time/growth after *t_min_* is observed. First, we assessed the effectiveness of beginning the second strike when the cell population has grown either 10%, 50%, or 150% larger than its minimum size following application of the first strike. This generally results in a high extinction probability (Figure 4E), but this probability decreases from >90% to ∼80% as we increase the amount of growth that we allow to occur between *t_min_* and the beginning of the second strike from 10% to 150%); the variance in outcomes also increases with the amount of lag time post-*t_min_*. Similarly, applying the second strike after a specific number of cycles (i.e. 2, 3, or 4 cycles) after *t_min_* again yields generally good results (Figure 4F), although again efficacy decreases and outcome variability increases as the amount of post-*t_min_* lag time before the second strike increases. In summary, our results suggest that while there may be variation in outcomes from patient to patient, a second strike is most likely to be successful if it can be initiated at or shortly after *t_min_*.

### Combination therapy outperforms sequential therapy, except when cross-resistance is present

To compare the efficacy of combination therapy to optimally timed sequential therapy, we carried out simulations like those of the sequential therapy scenario above, but also included the possibility of combination therapy, wherein both drugs are applied simultaneously and at an equal dose until population rescue or extinction. We compared these simulations in the presence and absence of the possibility of mutations conferring cross-resistance to both drugs. Our results (Figure 5A) indicate that in the absence of cross-resistance combination therapy outperforms optimally timed sequential therapy for all combination drug doses above 0.4, with combination therapy failing to achieve extinction for any replicates when the dose of both drugs is 0.4. Furthermore, we note for all cases where the combination therapy dose is >0.40, the extinction probability does not increase substantially with increases in the combination drug dose, regardless of the mutation rate. Indeed, it is worth stressing that the combination therapy outperforms two-strike extinction therapy even though in most of these parameterizations the individual drugs used in the two-strike therapy kill cancer cells at a much higher rate than the combined effect of the drugs in the combination therapy. This implies that it is the much-reduced probability of independently evolving resistance to both drugs rather than an elevated rate of killing cancer cells that is responsible for the greater efficacy of the combination therapy approach under this model.

**Figure 5.**
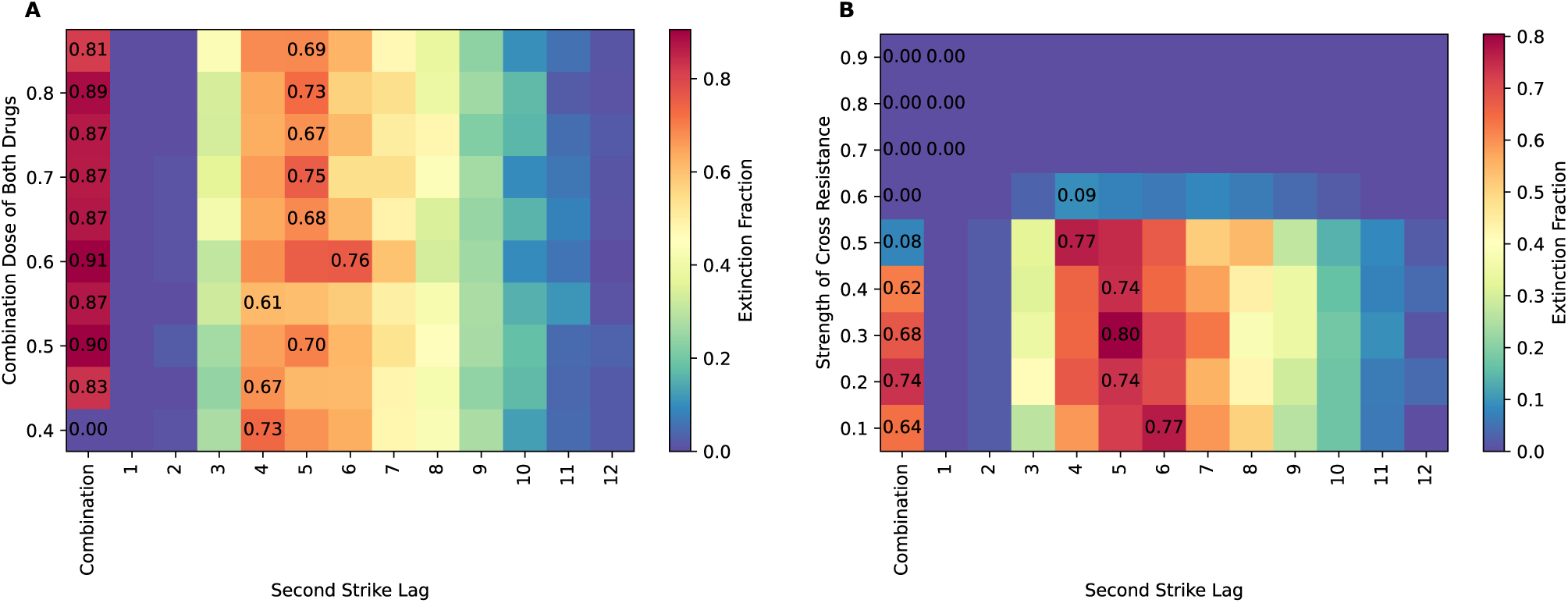
Comparison of efficacy between combination therapy and sequential therapy in the presence and absence of cross-resistance mutations shows that combination therapy is usually more effective except in cases of high cross resistance. A) Combination therapy and sequential therapy at a mutation rate of 5 × 10^-5^ in the absence of cross-resistance at various doses of combination drug. The sequential drug doses are fixed at 0.95 for both drugs. B) Combination therapy and sequential therapy at a mutation rate of 5 × 10^-5^ at various strengths of cross-resistance; the combination drug doses being fixed at 0.45 and the sequential drug doses are fixed at 0.95 for both drugs. The fraction of extinct replicates for the for the sequential protocol at the optimal time and for the combination protocol are indicated in black text. Combination therapy outperforms optimally timed sequential therapy at all doses of combination drugs, while sequential therapy outperforms combination therapy at the selected dose in the presence of cross mutations regardless of the strength of cross-resistance.

When cross-resistance mutations are present and the mutation rate is sufficiently high (Figure 5B), optimally timed sequential therapy outperforms combination therapy when the combination dose drugs are at 0.45, at all the tested strengths of cross-resistance. We also note that both therapies achieve extinction in only a small fraction of simulation replicates when the cross-resistance strength is high. This failure of treatment is more pronounced and occurs at even lower cross-resistance strengths when the mutation rate is 9 × 10^/1^ or higher. (See Figure S2 and Figure S3 for results at various mutation rates with and without cross-resistance). The lower probability of extinction under the sequential therapy at higher strengths of cross-resistance may be caused by cross-resistance mutations segregating at low frequency after the administration of the first drug. Such mutations are allowed to persist because the selective advantage of mutations conferring resistance to drug 1 over mutations conferring cross-resistance is diminished as the strength of cross resistance is increased, resulting in preexisting resistance to the second drug at the time of the second strike.

### Equal drug doses in combination therapy are optimal regardless of resistance mutation rates

Given the importance of the mutation rate to combination therapy efficacy, we tested whether there is an optimal ratio of drug doses during combination therapy if the two resistant mutations have different mutation rates. To achieve this, we carried out simulations involving only combination therapy, fixing the mutation rate for resistance to the first drug and varying the mutation rate of resistance to the second drug. As outlined in the Methods, we varied the drug dose ratio but kept their combined effect constant so that the likelihood of death for treatment-sensitive cells was the same across all simulations. Our results (Figure S4) indicate that regardless of the difference in drug mutation rates, the highest chance of inducing population extinction is achieved when the doses of the two drugs in the combination is 1:1. This makes sense as this scenario poses the greatest evolutionary challenge for the population in that both treatments need to be overcome to likely have significant rebound. Thus, our evolutionary rescue model suggests that it may be preferable to use a combination of treatments where each drug has a similar efficacy at killing cancer cells.

### Extinction therapy dynamics appear largely similar if treatment resistance is a quantitative trait

So far, our simulations have treated resistance as a binary trait. Evolutionary rescue has also been modelled as a quantitative trait (Gomulkiewicz R, Holt RD. 1995) where resistance involves multiple loci of modest effect. We tested whether the results above hold in scenarios where resistance is conferred by the cumulative effect of multiple mutations. The contribution of each mutation to resistance was drawn from a lognormal distribution as described in the Methods. We find that an optimal lag time still exists in these simulations, and that it remains independent of the mutation rate for resistance (Figure S5A). However, the peaks of extinction probability around the optimal lag times are not as sharp as those in binary resistance simulations (compare to Figure 1).

We also examined the impact of varying drug doses under our under this quantitative trait model, and draw similar conclusions as under our binary trait model: optimal lag times are independent of mutation rates regardless of which drug has a lower dose, optimal lag times remain unchanged when drug 2 has the smaller dose, and optimal lag times are higher and extinction probabilities are lower when drug 1 has the smaller dose (Figure S5B, C, and D).

## DISCUSSION

We performed forward-in-time agent-based population genetic simulations to evaluate the effectiveness of several different treatment strategies under an evolutionary rescue model. Under a sequential two-strike “extinction therapy” regime, we showed that the timing of the switch from the first drug to the second has a marked effect on the probability of the cell population going extinct (Figure 2). We also found that using the more effective drug first and the less effective drug second leads to substantially higher extinction probabilities than vice-versa (Figure 3). Furthermore, our simulations show that combination therapy forms a formidable barrier to the evolution of resistance when no mutations conferring cross-resistance exist, outperforming even optimally timed sequential therapy involving much stronger drugs than those used in the combination therapy (Figure 5A). This implies that combination therapies with lower doses may be preferable to sequential therapy using drugs that achieve a higher kill rate, because the probability of adapting to *each* drug in combination is very low so long as resistance mechanisms are independent—this result is consistent with insights gained from studies of pest control and antimicrobial treatment strategies (Pimentel and Burgess 1985; Angst et al. 2021). We note that these results are in line with another potential benefit of deploying combinations of drugs that attack different targets in a tumor cell: the time and cost of developing and evaluating new treatments could be significantly reduced by constructing a cocktail of existing pharmaceuticals rather than designing new drugs (Mokhtari et al. 2017).

On the other hand, even drug combinations that target distinct pathways of a cancer cell can often be overcome through a single mechanism that produces multidrug resistance such as elevated xenobiotic metabolism, increased growth factor production, etc. (Luqmani 2005; Bukowski et al. 2020). Our simulations show that when mutations that confer sufficient levels of cross-resistance can occur, combination therapy can yield worse outcomes than sequential therapy (Figure 5B). The superior effectiveness of two-strike extinction therapy in this scenario is because, in situations where cross-resistant mutations may be present but do not produce the same level of resistance as mutations conferring resistance to a specific drug, the two-strike therapy will cause single-drug resistance mutations to outcompete cross-resistant mutations, leaving the population more vulnerable to the next drug in the sequence. However, as the degree of resistance conferred by cross-resistance mutations approaches that of the single-drug resistance mutations, the advantage of sequential therapy is erased and both approaches fail with high probability. The failure of the two-strike approach here is most likely a result of the persistence of segregating cross-resistant variants following the application of the first drug which quickly render the second strike ineffective. We also show that with combination therapies it is optimal to use drug dosing where each drug kills cancer cells at the same rate, regardless of whether resistance to one drug arises at a higher rate than the other. Overall, our results support the need for developing drug combinations that are recalcitrant to the evolution of multidrug resistance (Pritchard et al. 2012), and suggest that when cross-resistance cannot be avoided, two-strike extinction therapy strategies may yield better outcomes.

We used *in silico* patient trials to investigate the evolutionary dynamics of two-strike extinction therapy in greater detail. We found that for many patients the optimal time to switch from the first to the second drug was at or near *t_min_*, the time at which the cancer cell population reaches its minimum and begins to rebound. Yet, the window of opportunity to switch therapy following *t_min_* may differ among patients. We do find that, if one switches treatments shortly after *t_min_*, the probability of extinction is often near its maximum (Figure 4). However, we note that because the absolute amount of growth occurring immediately after *t_min_* is relatively minimal (Figure 1 and Figure 4), determining *t_min_* may not be feasible in practice until well after it has occurred. Thus, if one waits until observable progression to initiate the second strike it may already be too late. We therefore argue that efforts to accurately predict *t_min_* for a given patient based on a limited number of scans/assays may prove to be a better route toward improving patient outcomes.

Our *in silico* trials show how adaptation to a treatment removes genetic diversity that could confer resistance to a second treatment. It is for this reason that in those patients for whom mutations conferring resistance to drug 2 were present at *t_min_*, we observed that the probability of extinction was very low if the second strike began at *t_min_*, but much higher if the duration of the first strike was slightly extended in order to remove these drug 2 resistance mutations. This illustrates that the success of two-strike extinction therapy shares some similarity to the strategy of imposing an evolutionary “double bind,” wherein resistance to one treatment confers susceptibility to another (Gatenby, Brown, et al. 2009; Basanta et al. 2012) although in our case this is a side effect of the reduction in tumor heterogeneity caused by selection rather than a direct phenotypic cost of resistance. In other words, the benefit of the first treatment is not only to shrink the size of the cancer cell population, but also to reduce the amount of diversity that could allow the population to immediately adapt to the second strike (Messer and Petrov 2013). If treatments are switched too early, mutations conferring resistance to the second strike may remain present in the cell population, while waiting too long allows the population to rebound and allows mutations conferring resistance to the second strike to randomly (re-)emerge in the population. Our results also imply that the strategy of repeated changes to the selective environment (i.e. treating a patient with a series of drugs used one after another in sequence) can further increase the probability of extinction. This is consistent with previous theoretical studies that have showed that frequent or continual environmental change can drive even initially very robust and adaptable populations to extinction (Bürger and Lynch 1995; Orive et al. 2019). Indeed, if environmental stochasticity is present in addition to a major environmental shift (i.e. treatment), extinction probability is further increased (e.g. (Xu et al. 2023)), a fact that future treatment strategies may be able to exploit.

We note that our conclusions regarding the optimal timing of the second strike are largely in agreement with those of a recent study by Patil et al. (2023) who constructed an analytical model of evolutionary rescue under a similar two-strike extinction therapy scenario as that examined here. Specifically, Patil et al. also found that the nadir of the population after the first strike is usually the optimal point to switch therapies, and that striking slightly after the nadir still yields good results. However, Patil et al. found that including a cost of resistance induces a more complex relationship where, if the second strike begins before the nadir, the extinction probability remains high for a longer duration when the first drug’s dose is intermediate and the second drug’s dose is high. This is because the population decline during the first strike is slower if the drug dose is not too high, and this allows more time for selection against cells resistant to the second drug, as these cells are experiencing the cost of resistance but no selective benefit. This result underscores the potential for resistance costs to qualitatively affect evolutionary dynamics during extinction therapy. The impact of the presence and strength of costs of resistance should thus be considered in future studies of not only the timing of two-strike extinction therapy but its relative efficacy vis-à-vis combination therapy.

Finally, the genetic architecture of resistance appears to have only modest implications for the effectiveness of sequential therapy. We observed similar dynamics whether treatment resistance is a binary trait or a quantitative trait where the degree of resistance is the sum of the effects of multiple mutations. However we note that in the latter scenario extinction probabilities display broader plateaus around the optimal lag times (Figure S5A). This may suggest that, all else being equal, sequential therapy involving treatments that exert highly polygenic selection on cancer populations may require less precision in the timing of the second strike, whereas treatments for which resistance is governed by a single locus may in some cases yield poor outcomes if the timing of the second strike misses the optimum by a relatively small period of time (Figure 2).

This study adds to the growing body of work showing that modeling the dynamics of evolutionary rescue can inform optimal treatment strategies and timings. The simulation approach that we adopted here is flexible. Our code can be easily adapted to model additional treatment strategies (e.g. adaptive therapy; (Gatenby, Silva, et al. 2009; Zhang et al. 2017; Zhang et al. 2022)) and to incorporate additional complexities into our model of cancer cell population evolution, adaptation, and extinction. Such features could include but are not limited to spatial heterogeneity and dynamics (Schmelz et al. 2021; Noble et al. 2022; Seferbekova et al. 2023), Allee effects (Boukal and Berec 2002; Konstorum et al. 2016; Brown et al. 2017; Gerlee et al. 2022), treatment refugia (Fu et al. 2015), non-genetic resistance mechanisms such as epigenetic responses (Bell and Gilan 2020) and transitions to drug-tolerant cell states (Sharma et al. 2010; Shaffer et al. 2017; Oren et al. 2021), and increasing rates of resistance mutations over time resulting from increased genomic instability (Lukow et al. 2021). Our agent-based simulation model could be readily extended to investigate these phenomena as well. More broadly, our study demonstrates the potential of flexible forward population genetic simulations of evolutionary rescue scenarios for investigating the dynamics of diverse treatment/management strategies for cancer and other diseases/organisms where the emergence of resistance is a common occurrence (e.g. antibiotic resistance, pesticide resistance, etc).

## ACKNOWLEDGMENTS

The authors thank attendees of the Triangle Center for Evolutionary Medicine’s 2023 Cancer and Evolution Catalysis Meeting for feedback on a preliminary version of this project.

## FUNDING

DRS was supported by National Institutes of Health award R35GM138286, and AD was supported by the Royster Society of Fellows at the University of North Carolina at Chapel Hill. CDJ acknowledges the University Cancer Research Fund and the Lineberger Comprehensive Cancer Center Grant (P30CA016086) for support.

## DATA AVAILABILITY

All code necessary for conducting the simulations carried out in this study can be found at https://github.com/SchriderLab/cancer_sims.

## Notes

### Competing Interest Statement

The authors have declared no competing interest.

### Summary of Updates

We have clarified the text in numerous places and also cited some relevant literature that was omitted from our initial submission.

