## Supplementary Figures for "Evolutionary rescue model informs strategies for driving cancer cell populations to extinction"

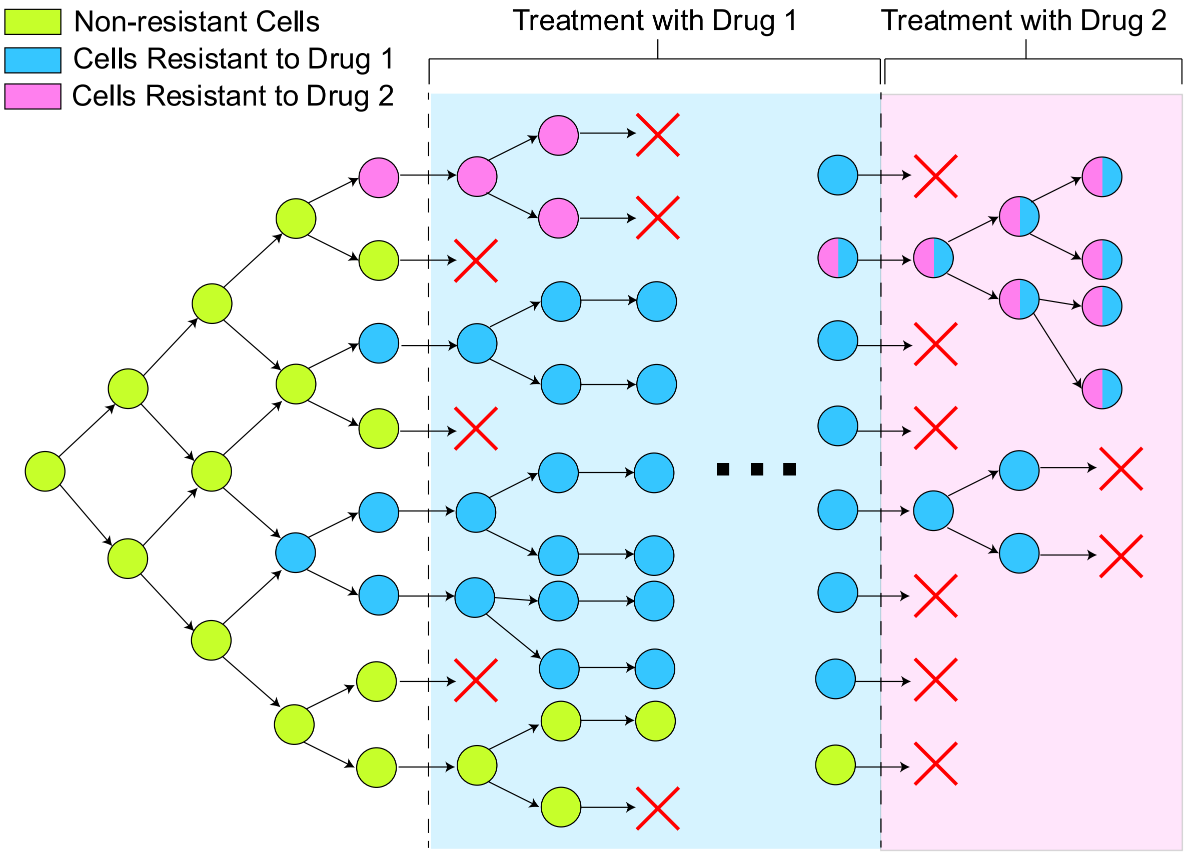


Figure S1. Conceptual simulation overview. The cancer cell population expands stochastically from a single non-resistance cell until reaching a population of within 10% of $2^{20}$ cells. The first strike is then applied continuously for a specific number of cycles (the lag time) before the second strike is then applied continuously until the population reaches extinction or rescue. Cells can accumulate mutations that confer resistance to each drug at any point during the simulation.

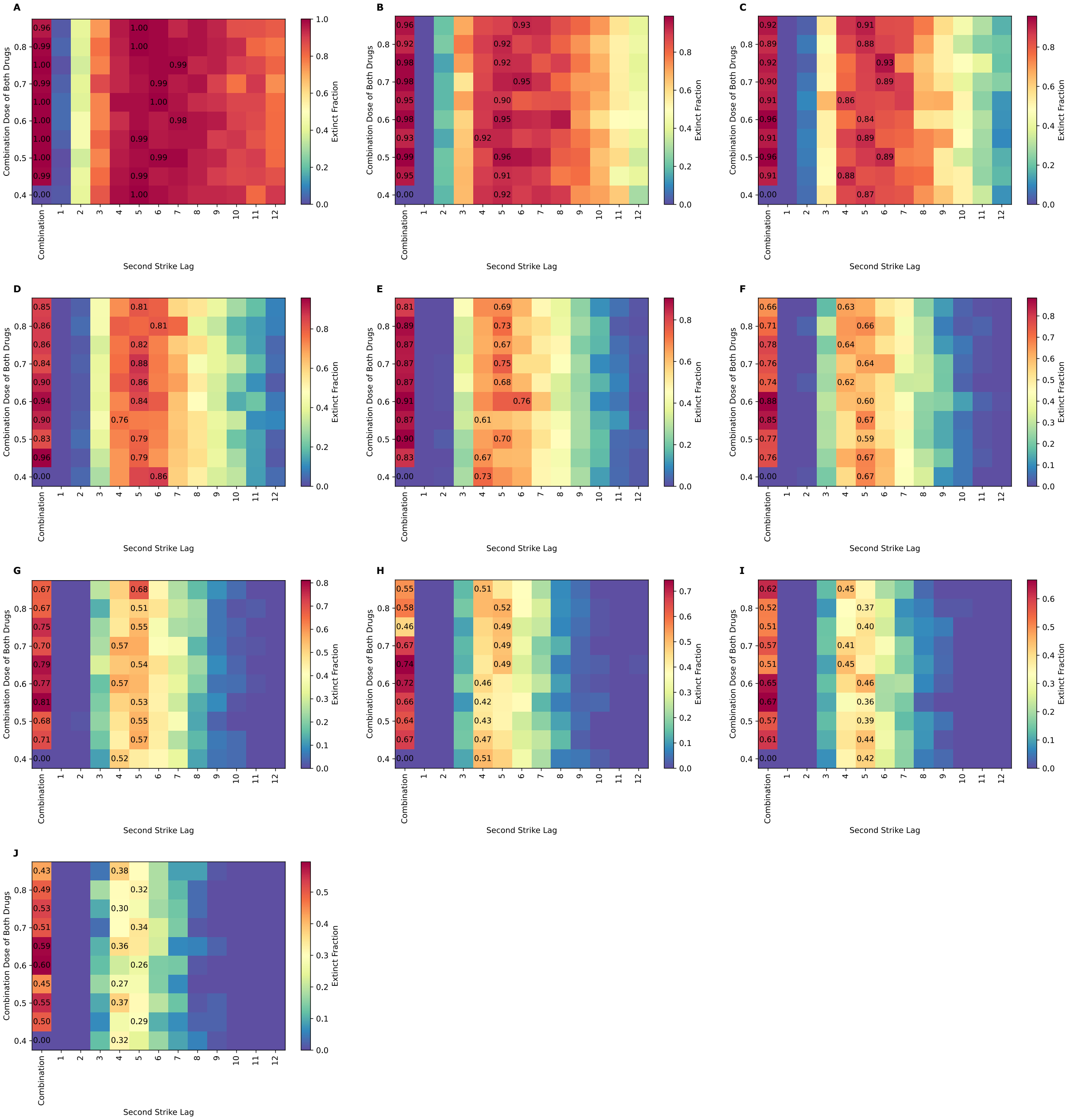


Figure S2. Comparison of combination therapy and sequential two-strike therapy protocols across various mutation rates and drug combination doses in the absence of cross-resistance mutations. Black text indicates the fraction of extinct replicates for combination therapy and at the optimal second-strike lag for sequential therapy for each drug combination dose.  The dose of both drugs in the sequential therapy scenarios is fixed at 0.90. Each panel represents a different mutation rate, with panel (A) representing a mutation rate of $10^{-5}$, increasing by $10^{-5}$ for each panel until reaching $10^{-6}$ in panel (J). Combination therapy outperforms optimally timed sequential therapy across all parameter combinations.

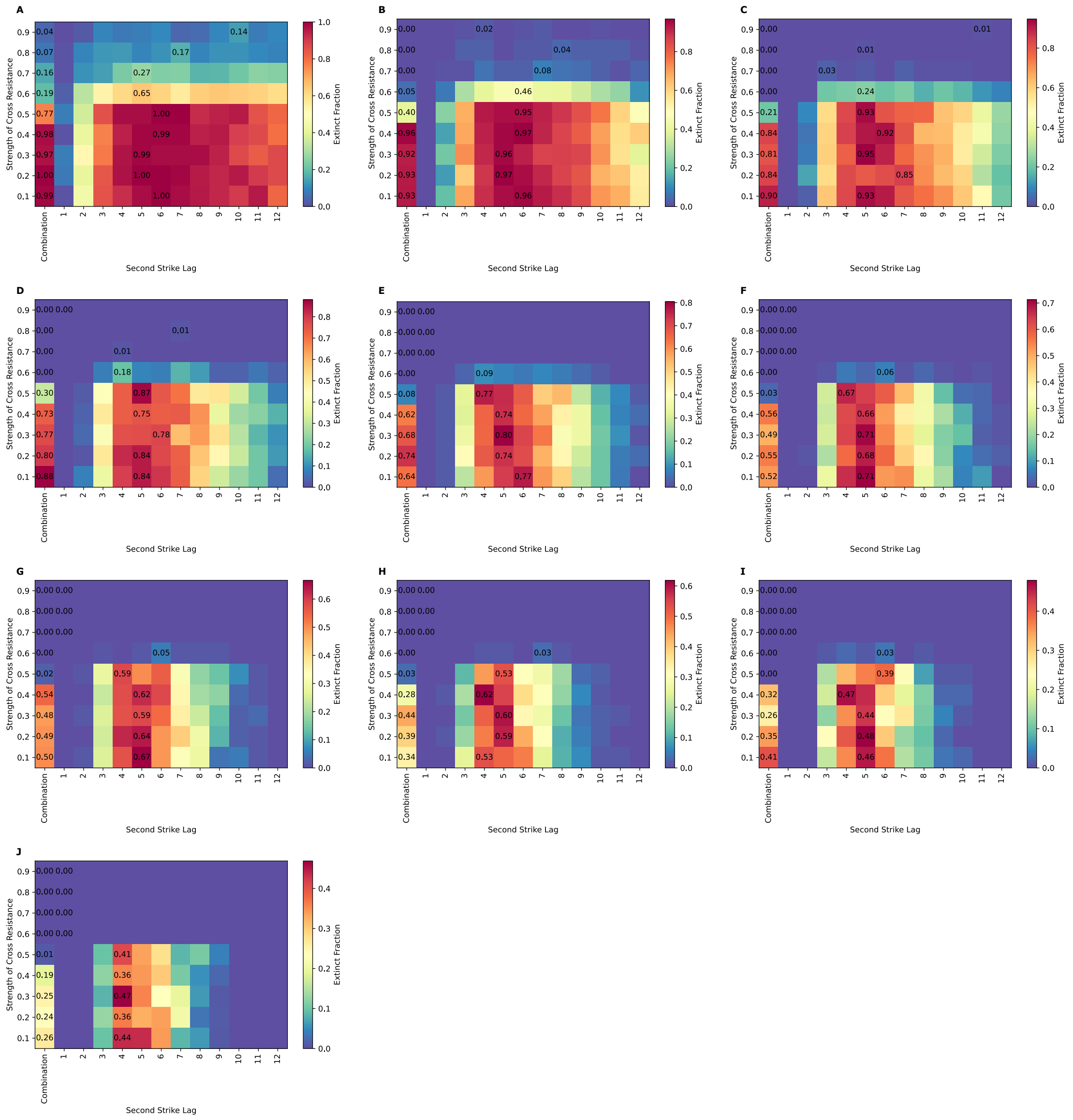


Figure S3. Comparison of combination therapy and sequential two-strike therapy protocols resistance in the presence of cross-resistance mutations across various mutation rates and strengths of cross-resistance. Black text indicates the fraction of extinct replicates for combination therapy and at the optimal second-strike lag for sequential therapy for each strength of cross-resistance. The dose of both drugs in the combination protocol is fixed at 0.45, while the dose for both drugs in the sequential therapy is fixed at 0.9. Each panel represents a different mutation rate, with panel (A) representing a mutation rate of $10^{-5}$, increasing by $10^{-5}$ for each panel until reaching $10^{-6}$ in panel (J). With the introduction of cross-resistance, sequential therapy outperforms combination therapy at the doses chosen, even at low levels of cross-resistance.

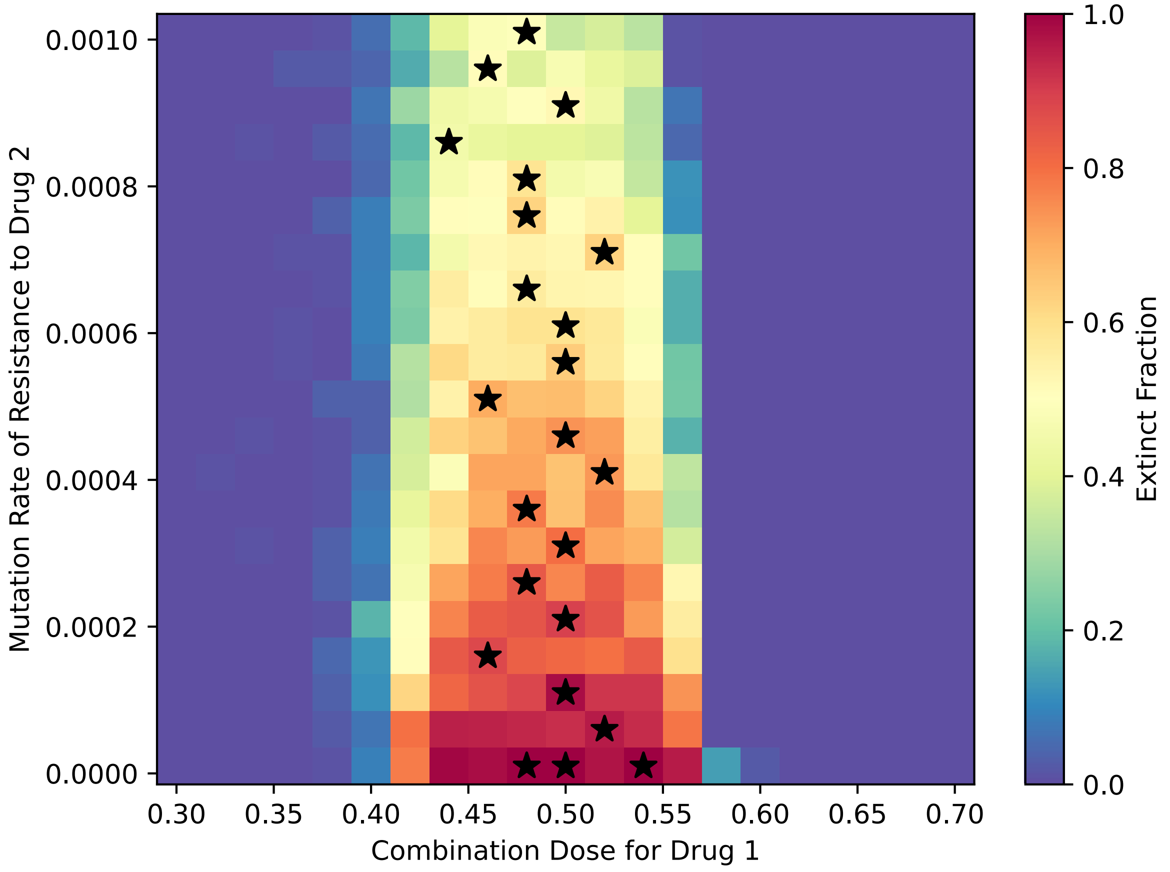


Figure S4. Fraction of extinct replicates are various doses of the first drug and rate of mutation of the second drug in a combination therapy scenario. The mutation rate for the first drug is fixed at $10^{-5}$, and the dose for the second drug is always chosen so that the probability of survival for non-resistant cells is the same. An equal or close-to-equal dose of both drugs is the optimal choice regardless of the mutation rate of resistance to the second drug. Black stars are drawn for each row at the combination dose with the highest extinct fraction, with multiple stars per row in case of a tie.

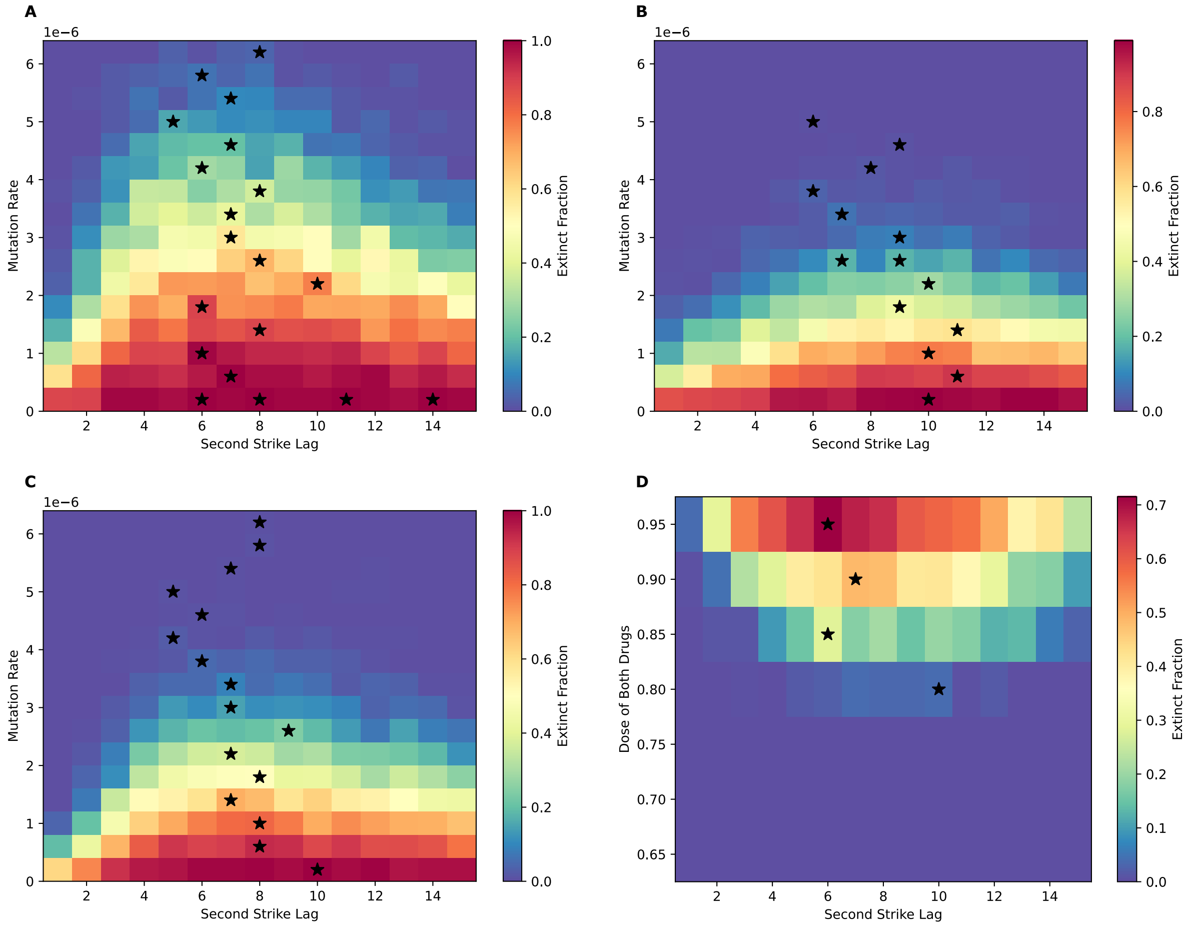


Figure S5. Results of simulations where resistance is considered a quantitative trait rather than a binary one, with 10 mutations of one type on average conferring full resistance in a cell to the corresponding drug. (A) The effect of the mutation rate and the second strike lag on the fraction of extinct replicates (extinct fraction), with the dose of both drugs fixed at 0.75. (B) The effect of the mutation rate and the second strike lag on the extinct fraction when drug 1 has a dose of 0.65 and drug 2 has a dose of 0.75. (C) The effect of the mutation rate and the second strike lag on the extinct fraction when drug 1 has a dose of 0.75 and drug 2 has a dose of 0.65. (E) The effect of the dose of both drugs and the second strike lag on the extinct fraction when the mutation rate is fixed at $3.2 \times 10^{\wedge}(-6)$. In general, conclusions from the corresponding binary trait resistance simulations hold, but the peaks of extinction probability around the optimal lag times are broader in the case of resistance as a quantitative trait.
